# Scaling deep phylogenetic embedding to ultra-large reference trees: a tree-aware ensemble approach

**DOI:** 10.1101/2023.03.27.534201

**Authors:** Yueyu Jiang, Daniel McDonald, Rob Knight, Siavash Mirarab

**Affiliations:** Electrical and Computer Engineering Department, University of California, San Diego, 9500 Gilman Drive, 92093, CA, USA; Pediatrics Department, University of California San Diego, 9500 Gilman Drive, 92093, CA, USA; Center for Microbiome Innovation, Jacobs School of Engineering, University of California San Diego, 9500 Gilman Drive, 92093, CA, USA

**Keywords:** Phylogenetic placement, Deep learning, Divide-and-conquer, Greengenes

## Abstract

Phylogenetic placement of a query sequence on a backbone tree is increasingly used across biomedical sciences to identify the content of a sample from its DNA content. The accuracy of such analyses depends on the density of the backbone tree, making it crucial that placement methods scale to very large trees. Moreover, a new paradigm has been recently proposed to place sequences on the species tree using single-gene data. The goal is to better characterize the samples and to enable combined analyses of marker-gene (e.g., 16S rRNA gene amplicon) and genome-wide data. The recent method DEPP enables performing such analyses using metric learning. However, metric learning is hampered by a need to compute and save a quadratically growing matrix of pairwise distances during training. Thus, DEPP (or any distance-based method) does not scale to more than roughly ten thousand species, a problem that we faced when trying to use our recently released Greengenes2 (GG2) reference tree containing 331,270 species. Scalability problems can be addressed in phylogenetics using divide-and-conquer. However, applying divide-and-conquer to data-hungry machine learning methods needs nuance. This paper explores divide-and-conquer for training ensembles of DEPP models, culminating in a method called C-DEPP that uses carefully crafted techniques to enable quasi-linear scaling while maintaining accuracy. C-DEPP enables placing twenty million 16S fragments on the GG2 reference tree in 41 hours of computation.

## Introduction

The past decade has brought much focus on phylogenetic placement of a query taxon on a backbone tree (Matsen et al., 2010; Berger et al., 2011; Mirarab et al., 2012; Zheng et al., 2018; Barbera et al., 2019; Linard et al., 2019; Rabiee and Mirarab, 2020b; Balaban et al., 2020; Turakhia et al., 2021; Wedell et al., 2022a). The reason behind this increased popularity is that placement can identify the contents of a biological sample, a problem that is enormously consequential in many downstream applications. Placement is used extensively in microbiome analyses (Matsen, 2015; Janssen et al., 2018) and tracking of epidemics (Turakhia et al., 2021). What placement offers, in lieu of the *de novo* reconstruction, is scalability: since placement processes queries independently, it scales linearly with the number of queries and enables analyzing millions of queries. This focus on scalability, however, should not come at the expense of accuracy.

A main lesson learned in analyses using existing tools, one that should not be surprising, is that the accuracy of the placements and downstream analyses both depend on the density of the backbone tree (e.g., Nasko et al., 2018; McDonald et al., 2022). For example, Balaban et al. (2020, 2022) documented that subsampling a larger tree to create smaller backbone trees reduced accuracy for all methods tested. Most methods have reduced accuracy when the closest matches in the reference database differ substantially from the query. This observation has spurred the development of many reference sets (many using genome-wide data) that include tens to hundreds of thousands of taxa (e.g., Quast et al., 2012; Shi et al., 2019; Parks et al., 2018; Zhu et al., 2019; Asnicar et al., 2020; McDonald et al., 2022). These large databases include a fraction of available prokaryotic genomes and a tiny fraction of an estimated 10^12^ microbial species (Locey and Lennon, 2016).

While placement methods are naturally scalable with more queries, they do not always scale to large backbone trees. This lack of scalability can hamper the use of methods and has motivated the development of booster methods such as pplacer-XR (Wedell et al., 2021) and SCAMMP (Wedell et al., 2022a) that scale existing methods. These methods extensively rely on a divide-and-conquer strategy that is used by all placement methods to some degree. Some of the existing methods, such as APPLES-II (Balaban et al., 2022) and SCAMMP, have been successfully run with reference trees with many tens of thousands of leaves with reasonable running times.

What connects most traditional placement methods is that they use some model of sequence evolution to place a query on a tree that has generated the sequences. A new paradigm some of us recently proposed (Jiang et al., 2022a) is to use the species tree as the backbone while sequence data come from a single or a handful of genes. While this so-called discordant placement is conceptually less appealing than the traditional approach, it is very useful in practice. The goal of sample identification is to find species identities, not genes; discordant placement makes that conflict explicit. Also, by allowing updates of a species tree using single genes, it provides a path for combining two types of data historically analyzed separately: genome-wide (shotgun) metagenomic data and 16S amplicon-based data. If we can place 16S data on the species tree, we can jointly analyze 16S and genome-wide data. Jiang et al. (2022a) demonstrated that this goal is achievable with decent accuracy. In particular, they proposed a metric learning framework for phylogenetic placement using deep neural networks. The method proposed, called DEPP, trains a model based on backbone tree and alignment and uses the trained model to place new queries. The trained model is an embedder that maps sequences into R*d* space such that Euclidean distances in the embedding space match the patristic distances on the backbone tree (see Jiang et al. (2022b) for an extension to hyperbolic spaces).

While DEPP was successfully tested on backbone trees with roughly 10,000 species, it has fundamental scalability issues that need addressing. DEPP requires calculating an *O*(*n*^2^) distance matrix for training on a backbone tree with *n* leaves. For datasets with *n* ≫ 10^4^, saving the matrix (not to mention calculating it) is impossible on most machines. The alternative, to compute distances on the fly without saving them in the memory, is too time-consuming and makes the training process too slow. Thus, the current DEPP is limited to roughly 10,000 species in its training dataset, making it unable to take advantage of the modern ultra-large reference sets.

This limitation is not just theoretical. In a recent effort, McDonald et al. (2022) built a new version of the widely used Greengenes (DeSantis et al., 2006) reference dataset (GG2) complete with a tree with 331,270 tips, including both genomes and 16S sequences. Because this tree is (partially) a species tree, placing 16S rRNA amplicon sequences on the tree is best done using DEPP. However, DEPP cannot handle such a large backbone tree. To enable using GG2, we needed to develop a set of techniques that enable DEPP to scale to much larger backbone trees. These methods use divide-and-conquer to make the training time and memory grow quasi-linearly with the size of the backbone. However, as we show, much care is needed to retain the accuracy of the original method, mainly because of the trade-off between generalizability and precision during model training. This paper reports various ways of scaling DEPP, culminating in a method called Clustered-DEPP (C-DEPP), which is used by Greengenes2 (GG2). Besides showing the scalability and accuracy of C-DEPP, our results shed light on more basic questions about the best ways to perform phylogenetic divide-and-conquer in the machine-learning context.

## Methods

### Problem statement

#### Phylogenetic Placement and updates

The phylogenetic placement problem seeks to determine the optimal position of a query species on a backbone tree *T* consisting of species *n* accompanied by corresponding sequences. We also study the related tree-update problem: Given the backbone tree and the corresponding sequences, extend the tree to include the new species. Unlike placement, the resulting tree in the tree update task is a fully-resolved tree that elucidates the relationships between queries. Jiang et al. (2022a) introduced the concept of *discordance* phylogenetic placement where the backbone tree is not solely or exclusively inferred from the sequences used for placement. For downstream applications, placing on the species tree is the ultimate goal, even when data from a single gene is available. In this paper, we focus on discordance placement and update.

### Training/Testing Datasets

We will focus on two datasets for benchmarking.

#### Biological data

The Web-of-Life (WoL) dataset, built by Zhu et al. (2019), contains 10,575 species and 381 marker genes. An ASTRAL tree constructed using the 381 marker genes is available with the branch length calculated using sites sampled from the marker genes. Here, we use the 10 genes examined by Jiang et al. (2022b) as well as the marker 16S gene. Because the dataset is at the limit of what DEPP can analyze, using this dataset, we can compare the effect of various scaling strategies to the baseline method trained on the entire dataset. To allow fair comparisons, we use the same set of queries used by Jiang et al. (2022b) for phylogenetic placement and tree update. For placement, 5% of the species of each gene are randomly selected as the queries and removed from the backbone. For the tree update, we have two replicates. In each one, 100 random clades in the species tree ranging in size from 5 to 10 species were selected and pruned from the tree, with the remaining species serving as the reference. In total, across 11 genes, we have 5,038 queries for placement and 12,248 queries for tree update.

#### Simulated data

Similar to Jiang et al. (2022a), we generated a dataset comprising a species tree with 64,000 species and 100 genes undergoing extensive horizontal gene transfer (HGT) and some incomplete lineage sorting (ILS) using Simphy (Mallo et al., 2016). The branch lengths of the species tree were estimated under the GTR model using sequences from the 100 genes, with each gene providing 100 randomly selected sites. Here, we used only the first 5 genes for the placement experiments. For identical sequences, a random species was retained and all others were removed. This step resulted in the removal of 505 to 1,935 sequences among the five genes. Then, 5% of the species were randomly removed from the species tree as queries, resulting in 15,694 queries across all the genes.

### Background: DEPP

DEPP uses machine learning to estimate evolutionary distances between sequences. Unlike traditional methods, which rely on predefined evolutionary models, it learns to embed sequences in the *d*-dimensional space such that the pairwise distances of the embeddings approximate the tree distances, either in Euclidean space or Hyperbolic space (Jiang et al., 2022b). In the training phase, DEPP uses stochastic gradient descent to set the parameters of the model to minimize the cost function:

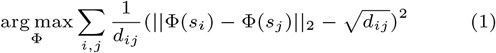

where *d*_*ij*_ are the backbone tree distances and Φ(*s*_*i*_) are the output embeddings generated by the DEPP model. The model is a neural network that consists of a single convolutional layer followed by a residual block, which comprises two convolutional layers with the input being added to the output. A final fully-connected layer is appended to generate the embeddings. Once the models are trained on the backbone tree and the backbone sequences, we can use them to place queries. To do so, we use the model to compute a distance vector and use this distance vector as the input to the distance-based phylogenetic placement method APPLES-2 (Balaban et al., 2022), which finds the optimal placement.

### C-DEPP: Scaling DEPP

To scale DEPP to *n* ≫ 10^4^, we use a divide- and-conquer strategy in our proposed method C-DEPP. To summarize (Figure 1), C-DEPP trains a separate model for each of several overlapping subtrees; for each query, C-DEPP uses a 2-level classifier to select one or more subtrees, computes distances using those subtrees, and uses these distances as input to APPLES-II, leaving the other distances blank. For this strategy to work optimally, many algorithmic tricks are needed. To motivate our final approach, below, we propose successively more advanced strategies and discuss the shortcomings of each. Since it is unclear how to evaluate these strategies theoretically, we resort to *empirical* evaluation to show that each additional strategy does contribute to better accuracy. We use two datasets mentioned earlier for benchmarking. The WoL dataset allows us to compare to normal DEPP while the simulated dataset allows examining the effects of having a very large training set and abundant HGT.

**Fig. 1.**
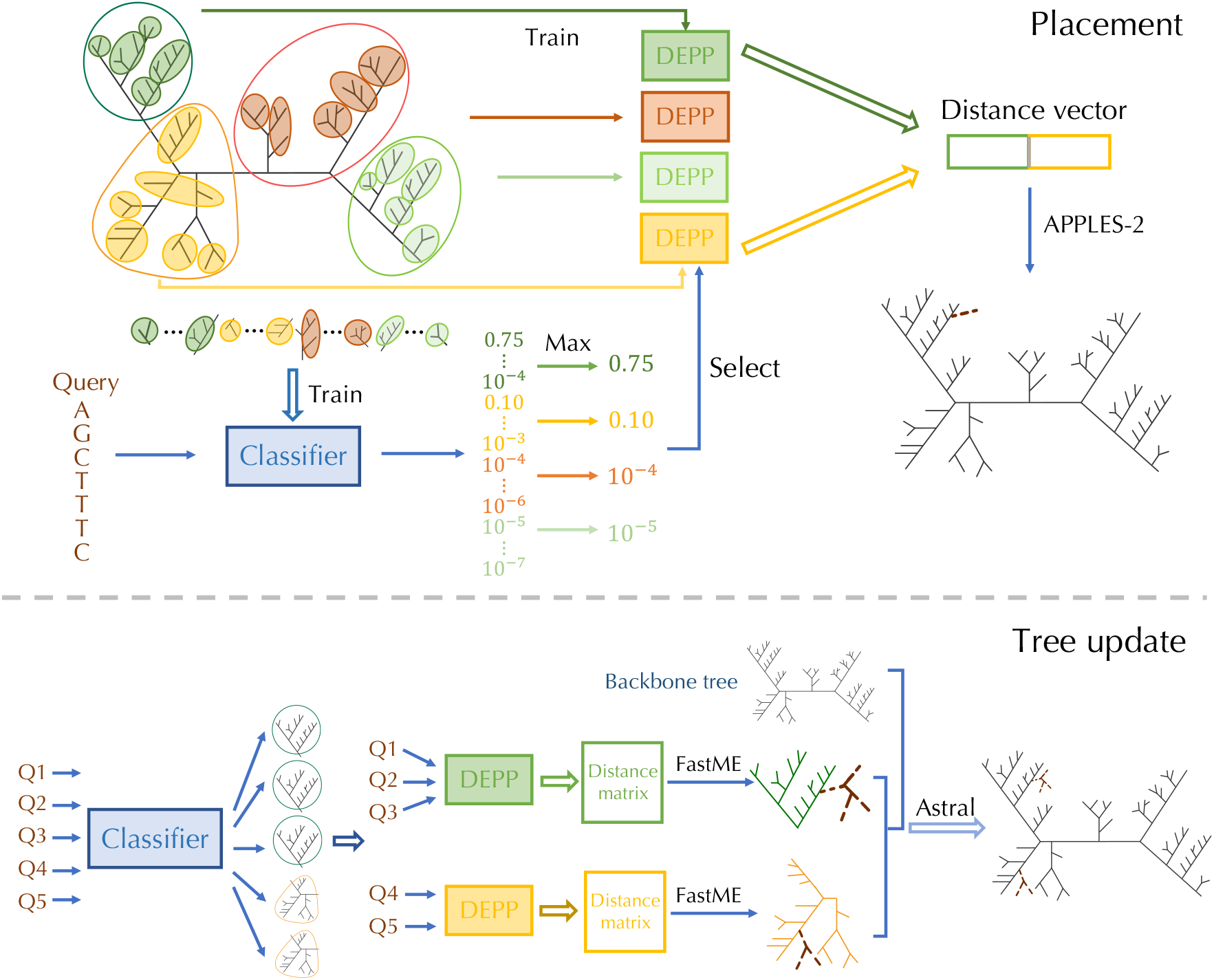
Pipeline of C-DEPP for phylogenetic placement and tree update. The tree is divided into multiple 1st-level groups and a separate neural network is trained on each of these subsets. Note that representatives from one group are added to the other ones during training (not shown here). Each 1st level subset is further divided into smaller subsets. A classifier is trained to classify a query to one of these smaller subsets probabilistically. At the query time, probabilities from small subsets are summarized (max) to 1st-level subsets, and the corresponding models are used to calculate distances. APPLES-II is used to place on the backbone, using only distances from chosen models. Updates happen similarly, but using FastME to update subsets and using ASTRAL (as a supertree method) to combine the subset trees and the original backbone.

### Subsampling

The most obvious option for scaling is to simply train the model on a subset of species available in the tree. Such subsampling would still allow placement on the full tree as the model can embed the unused backbones as well. However, the accuracy of deep learning models is known to depend on the size of training sets. Moreover, taxon sampling is crucial to phylogenetic accuracy (Zwickl and Hillis, 2002) and phylogenetic placement (Balaban et al., 2022). Thus, we expect subsampling to reduce the accuracy. On the biological WoL dataset, reducing sampling by ten folds increases the average error by around 50% (Fig. 2).

**Fig. 2.**
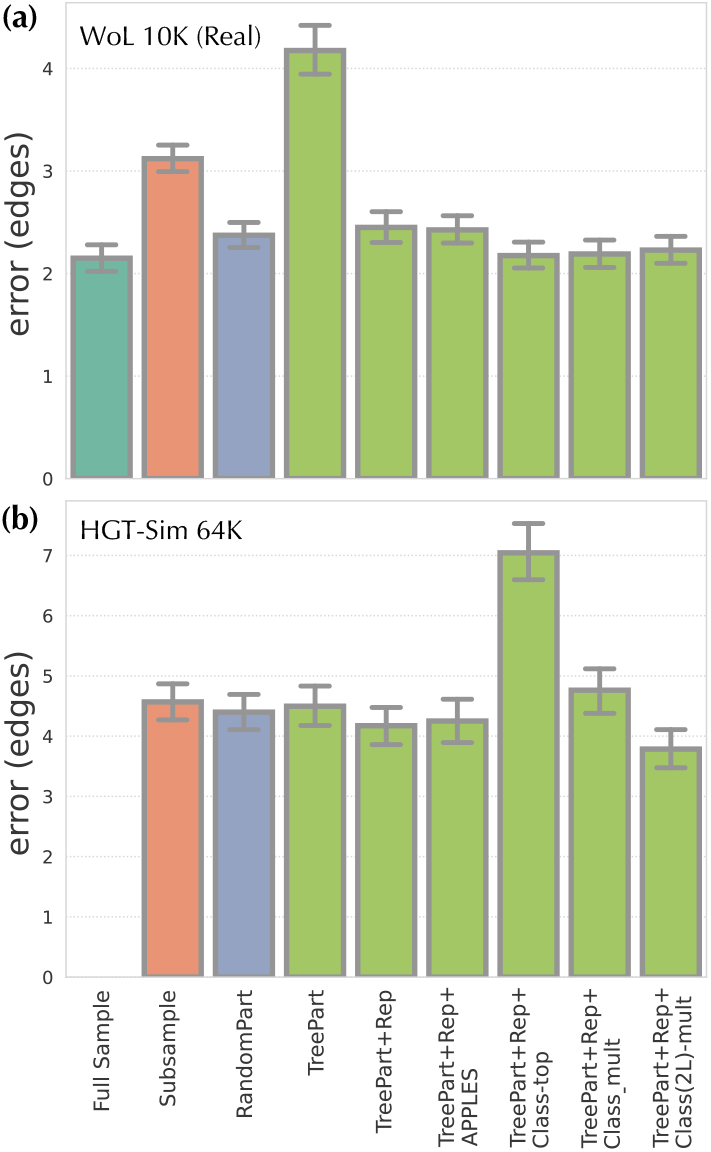
Performance of different strategies for scaling DEPP to ultra-large trees, tested on (a) WoL dataset, averaged over the 11 marker genes and (b) an example gene of the simulated dataset with *n* = 64000 leaves. We show the mean and standard error of the placement error. The full dataset can only be used for training on the WoL dataset. We compare it with strategies of subsampling the dataset to 2,500 (a) and 3,500 (b) sequences (similar to partition sizes used in the remaining methods), partitioning the dataset randomly into 3 (a) and 17 (b) subsets, and C-DEPP. For C-DEPP, features are added sequentially; TreePart: tree-based clustering; Rep: augment clusters with representatives from other groups; Class: use a classifier to select one (-one) or multiple (-mult) clusters; Class(2L): 2-level classification scheme. TreePart+Rep+Class(2L)-mult is the final version used elsewhere.

### Random Partitioning

An alternative is to partition the data and train a separate encoder on each subset. The resulting *ensemble* model allows calculating distances to each backbone using the associated model. Implementing this ensemble model using a random partitioning of data is far better than subsampling (compare Subsample and RandomPart in Figure 2) and comes close to the accuracy of the original model built on the full dataset. Nevertheless, it is less accurate and leads to the question: does more careful partitioning help?

### Tree-based partitioning

Instead of random partitioning, we can use the reference tree to create subtrees that are more evolutionary homogeneous. Such an approach creates a mixture of local experts, a well-established concept in machine learning (Jacobs et al., 1991) and used previously for phylogenetic placement (Mirarab et al., 2012). While many tree decomposition methods are available (e.g., Liu et al., 2011), we base our approach on the following criterion: divide the tree into subsets that are at most of the size *m* while minimizing the number of clusters. The optimal solution can be found in time linear in the size of the tree *n*, as implemented by the TreeCluster (Balaban et al., 2019) method. Thus, the tree is divided into subtrees *T* = {*t*_1_ … *t*_*c*_}, each with at most *m* species. We fix *m* (1500 by default) as *n* changes; thus, running time and memory both scale linearly with *n*.

Surprisingly, phylogenetic partitioning has a higher error than random partitioning (compare RandomPart and TreePart in Figure 2). This reduced accuracy may be due to the fact that each model is trained on a subtree without the ability to learn from the full range of possibilities in the sequence space. Thus, the sequence embedder trained on less diverse data is perhaps more precise but less generalizable (i.e., is overfit).

### Adding representatives (overlapping clusters)

To address the lack of generalizability, we design an approach that still uses the tree but creates overlapping subsets: Each subset includes *all* of sequences in one of the phylogenetic partitions created previously plus a selection of sequences from other subsets. More precisely, we create another set of subtrees 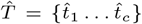 by adding *k* auxiliary species to each subtree in *T*. We set 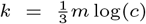 by default. Note that as we fix *m* when *n* grows, the size of the 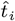 subtrees grow with *O*(log(*n*)) and the total running time and memory grow with *O*(*n* log^2^(*n*)). The auxiliary species added to *t*_*i*_ are those with the minimum distances to *any* species in *t*_*i*_. Since the subtrees are overlapping, the distance of a query to a reference species can be calculated using multiple models; when this happens, we simply take the median distance.

This overlapping partitioning approach performs better than random and tree-based partitioning alone (compare TreePart+Rep to TreePart and RandomPart in Figure 2). We believe the reason is that each model in the ensemble is both a local expert and also aware of the larger context (e.g., sequences outside the subtree). Thus, its embeddings are better than other “experts” for sequences belonging to that subset but it also is not ignorant of the rest of the space. Such hybrid approaches have been also used previously in machine learning (Peralta et al., 2019; Liao et al., 2019).

### Selecting the best model(s): classifiers

Instead of simply using all distances computed from all the models, would it be better to pick the best “expert” model and use only its distances? Note that APPLES-II allows missing distances, enabling us to place on the full tree using distances computed from a single model. Similar to methods such as SCAMPP and pplacer-XR (Wedell et al., 2022a, 2021), we can use an initial placement using APPLES+JC to pick a subset and use only the model trained on that subset for placement. This approach works no better or worse than giving APPLES-II the distances from all the models (compare TreePart+Rep+APPLES and TreePart+Rep in Figure 2). But can we do better than using an initial placement?

Deciding which cluster to use can itself be posed as a classification problem where the input is a sequence, and the output is a probability vector indicating the likelihood of the input sequence belonging to each subtree. Using an architecture very similar to DEPP, we designed such a classifier. The only difference compared to DEPP is that the number of embedding dimensions equals the number of partitions and the final output goes through a softmax layer that ensures the *L*_1_ norm of the output is 1 (i.e., can be interpreted as the probability of the partition). The loss function is the cross-entropy between the output probabilities and the ground truth (i.e., indicator function of the correct partition).

Using this classifier and simply picking the most likely cluster improves accuracy in the real dataset but dramatically reduces the accuracy in the simulated high-HGT dataset (TreePart+Rep+Class-Top vs TreePart+Rep in Figure 2). We then resorted to using multiple models when they all have a substantial likelihood. More precisely, we sort the models based on their likelihood and take each of the top 4 models if it has a likelihood at least 1/200 times the likelihood of the previously taken model (the threshold was picked arbitrarily and not optimized). This strategy substantially improves results (TreePart+Rep+Class-Multi in Figure 2) but still remains slightly worse than using all the models on the simulated dataset. We believe the reason is that the classifier needs to assign a sequence to very large groups, a task that may be difficult in the face of HGT among distantly related species. Recall that the backbone tree (used in partitioning) is the species tree and may not reflect the relationships among genes, making the high-level classification more challenging.

### 2-Level classifier

We propose a 2-level classification scheme. For the second level, each subtree in *T* (e.g., *t*_*i*_) is further split into smaller subtrees 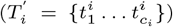 with a maximum of *m*’ leaves, where *m*’ is set to 30 by default. The classifier is then trained to select among the second-level subtrees 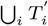. This classifier is built identically to what we described earlier for 1-level classifiers. To determine the subtree of a query, the classifier is used to calculate the likelihood 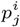 of the query belonging to each 2nd-level subtree *j* of 1st-level subtree *i*. Next, we define the score of the 1st-level subtree *t*_*i*_ to be 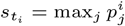 and assign the query to one or more 1st-level subtree(s) with the highest score. We keep assigning the query to up to four subtrees with the highest scores as long as the subset has a score that is at least 1*/*200 of the score of the previous subtree. Note that the 2nd-level classifiers are only used to select the 1st-level classes. Once the query is assigned to the 1st-level subtree(s), it is placed using distances calculated corresponding 1st-level model.

The 2-level classification strategy retains the same accuracy as the 1-level for the real dataset and substantially improves the accuracy for the simulated dataset (Figure 2). This version of the tool including all these features is the final version we will use throughout the rest of the paper as C-DEPP.

## Experimental Details

To evaluate the performance of C-DEPP, we compare our method with several alternative methods for both placement and update tasks.

For placement, we measure the number of edges between the placement and the correct placement on the tree. For the simulated dataset, we use the true species tree as the backbone, and thus, the correct placement is well-defined. On the real dataset, we take the position of the species in the original ASTRAL tree inferred from all 381 marker genes (before removing the queries) as the optimal position. We compare DEPP (only tested on the smaller real dataset) and C-DEPP to the following methods.

EPA-ng (Barbera et al., 2019) performs maximum-likelihood phylogenetic placement. We use RAxML-ng (Kozlov et al., 2019) to infer the parameters of the substitution models and the backbone tree under the GTR + Γ model.

EPA-ng-SCAMPP (Wedell et al., 2022a) is a method that enables EPA-ng to work on ultra-large trees by first finding a subtree and placing on that subtree using EPA-ng. We apply the default backbone tree size of 2000. Similar to EPA-ng, we use RAxML-ng to prepare the backbone parameters under the GTR + Γ model.

APPLES2+JC: We use APPLES-2 (Balaban et al., 2022) which uses Jukes-Cantor (JC) model to estimate distances. We use RAxML-ng under JC models to recalculate the branch length of the input backbone tree for this tool.

For tree updates, we measure both the Robinson and Foulds (1981) (RF) and quartet distance between the true/reference tree and the inferred updated tree. We compare these methods.

RAxML (Stamatakis, 2014) maximum-likelihood inference is used to update an existing tree; we use the backbone tree as a constraint to fix its topology. When multiple genes are available, we concatenate the sequences from all the genes.

JC+FastME. We first calculate the distances between all pairs of the sequences under the JC model and use the distance matrix as the input to the distance-based FastME (Lefort et al., 2015). When using more than one gene, we first calculated the distance matrix for each gene and then take the median of the distances for each pair across all genes to summarize the distance matrices. Distances among pairs of backbone species are fixed to patristic distances in the backbone tree to encourage FastME to keep the backbone relationships fixed.

DEPP+FastME is run using a pipeline similar to JC+FastME, using DEPP models to estimate the distances between sequences rather than the JC model.

C-DEPP+FastME first trains the models from the backbone and then performs three steps (fig. 1). First, we assign each query to the 1st-level subtree with the highest score *s*_*i*_; when multiple genes are available, we simply average the scores across genes. We calculate the distance matrices for all query and backbone species in each subtree using all genes. As in the previous methods, we take the median across genes and fix distances among backbone species to their patristic distance on the backbone tree. Whether we have single or multiple genes, we obtain a single distance matrix at the end for each subtree. We then re-infer each subtree using FastME given this distance matrix. To combine all the subtrees into a full tree, we give updated subtrees as well as the backbone tree as input to ASTER software (Zhang and Mirarab, 2022) without weighting. Note that, here, we use ASTER as a supertree method and not as a summary method combining gene trees.

## Results

### Simulated data

We start by evaluating C-DEPP on simulated data. Note that on this dataset, DEPP would require more than 250Gb of memory, and hence, we could not run it.

Comparing the remaining three methods, EPA-ng-SCAMPP has the best placement accuracy closely followed by C-DEPP (Fig. 3). Specifically, EPA-ng-SCAMPP has an average error of 2.78 edges over all five genes compared to 2.89 edges for C-DEPP. Both of the two methods are substantially more accurate than APPLES+JC with an average error of 3.73 edges. When evaluating the entire distribution of the error, the trend is generally consistent but a long tail of high errors is observed for all methods (Fig. S7). EPA-ng-SCAMPP finds placements at most three edges away from the correct placement 91.7% of the time, which is only 2.4% higher than C-DEPP and 11% higher than APPLES+JC.

**Fig. 3.**
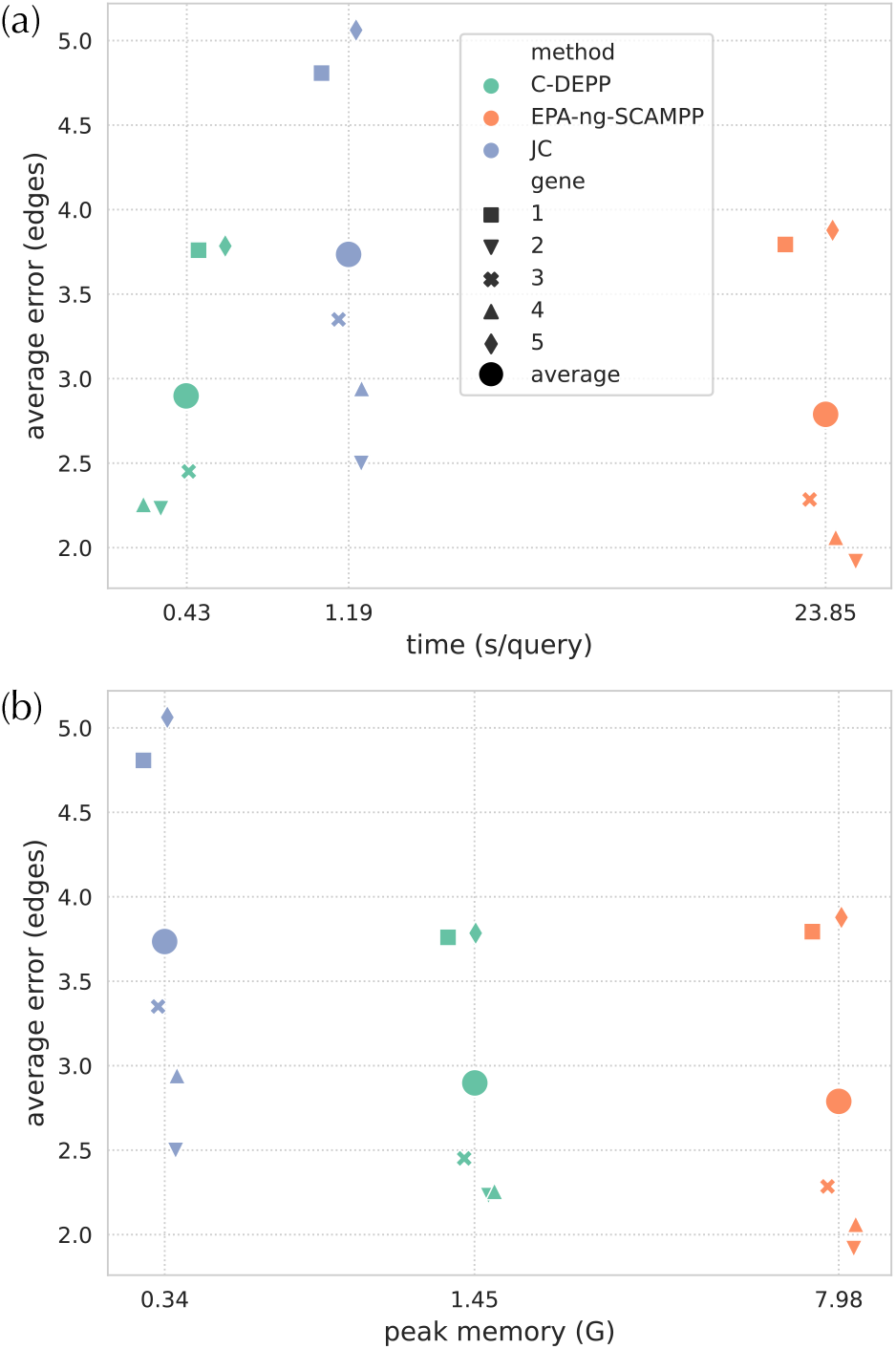
Results on HGT-Sim (64K) comparing average placement errors (y-axis) with running time (a) and peak memory (b). Larger dots are averages over all five replicates.

While slightly more accurate, EPA-ng-SCAMPP has much higher computational demands in terms of both running time and memory. When examining the running time, both C-DEPP and APPLES+JC are much faster than SCAMPP-EPA-ng (Fig. 3). Specifically, C-DEPP is 50 times faster than EPA-ng-SCAMPP, with a slightly higher error of 0.1 edges. Interestingly, C-DEPP is also faster than the less accurate method APPlES+JC. In terms of memory consumption, APPLES+JC is the most efficient requiring less than 0.5Gb, followed by C-DEPP which requires less than 1.5Gb; EPA-ng-SCAMPP required 8Gb on this dataset.

### Biological data

We now examine the results of WoL data.

#### Placement

Similar to the observation in the simulated data, the maximum-likelihood method EPA-ng outperforms the other alternatives in terms of placement accuracy (fig. 4). Averaged across the 11 genes, EPA-ng has the lowest error of 1.9 edges. Following EPA-ng, DEPP has the second lowest error of 2.1 edges. The performance C-DEPP is close to DEPP with a slightly higher average error of 2.2 edges. All these three methods are significantly more accurate than APPLES+JC whose average error is 3.1 edges.

**Fig. 4.**
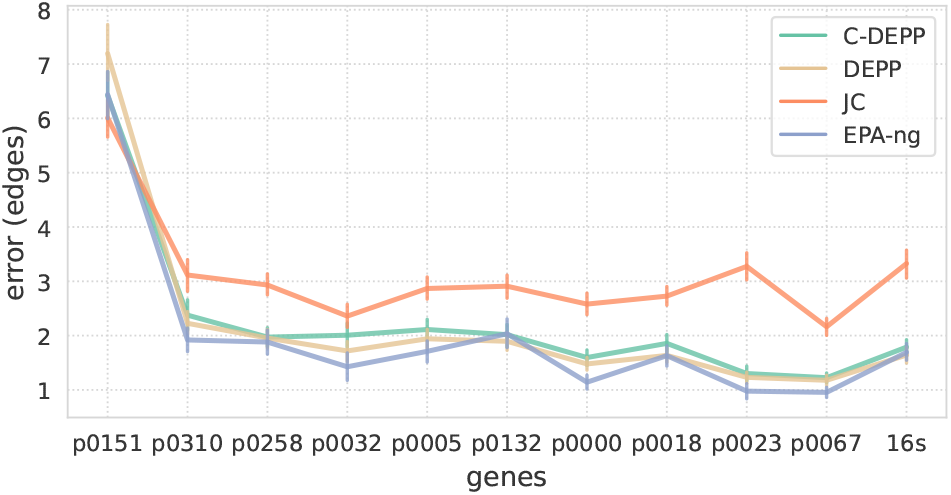
Placement error, showing mean and standard error of error, across 11 genes of the WoL dataset with *≈* 10^4^ species, where DEPP can be run.

While maintaining the high accuracy of DEPP almost intact, C-DEPP significantly reduces training time. For instance, training the DEPP model on 16S data from the WoL dataset using a Tesla V100-SXM2-32GB card takes 210 minutes. In contrast, the C-DEPP model requires only 1/3 of that time to train on the same data. In terms of testing, while C-DEPP has a longer running time than DEPP, it is more memory-efficient. For example, when placing 1000 16S sequences onto the backbone tree with 7400 leaves, C-DEPP has a peak memory usage of 0.67G (126 seconds running time), while DEPP requires a peak memory usage of 1.5G (26 seconds), reducing memory consumption by more than half compared to DEPP’s. Note that on datasets much larger than 7400 leaves, the memory requirements of

#### Tree update

When using a single gene, RAxML has a clear advantage over other methods with lower error measured by quartet distance or RF distance (fig. 5). However, C-DEPP has very similar accuracy to DEPP. When multiple genes are used, patterns gradually change. Errors drop rapidly for DEPP and C-DEPP but not the concatenation-based RAxML. For example, by using two genes, the average quartet distance of C-DEPP is 0.02 which is around 1/3 of its error with one gene. The quartet error quickly drops down to 0.006 with six genes. The pattern is similar (though less pronounced) when examining the RF distances. For example, the RF distances reduce by half from using a single gene to using six genes. In contrast, the error reduction is less pronounced for RAxML; the quartet error does not reduce notably in response to increasing the number of genes beyond two and can occasionally increase (e.g., from two genes to four genes). C-DEPP and DEPP start to outperform RAxML with six genes or more when measuring the quartet distance or with 10 genes when measuring the RF distance. The performance of JC+FastME is significantly worse than the other methods measured by quartet distance and is the worst or among the worst methods measured by RF distance. Finally, note that DEPP and C-DEPP have very similar accuracy. When the dataset is small enough to allow running both, there is no benefit in using C-DEPP on this dataset. However, for larger datasets, C-DEPP is the only option possible due to its size.

**Fig. 5.**
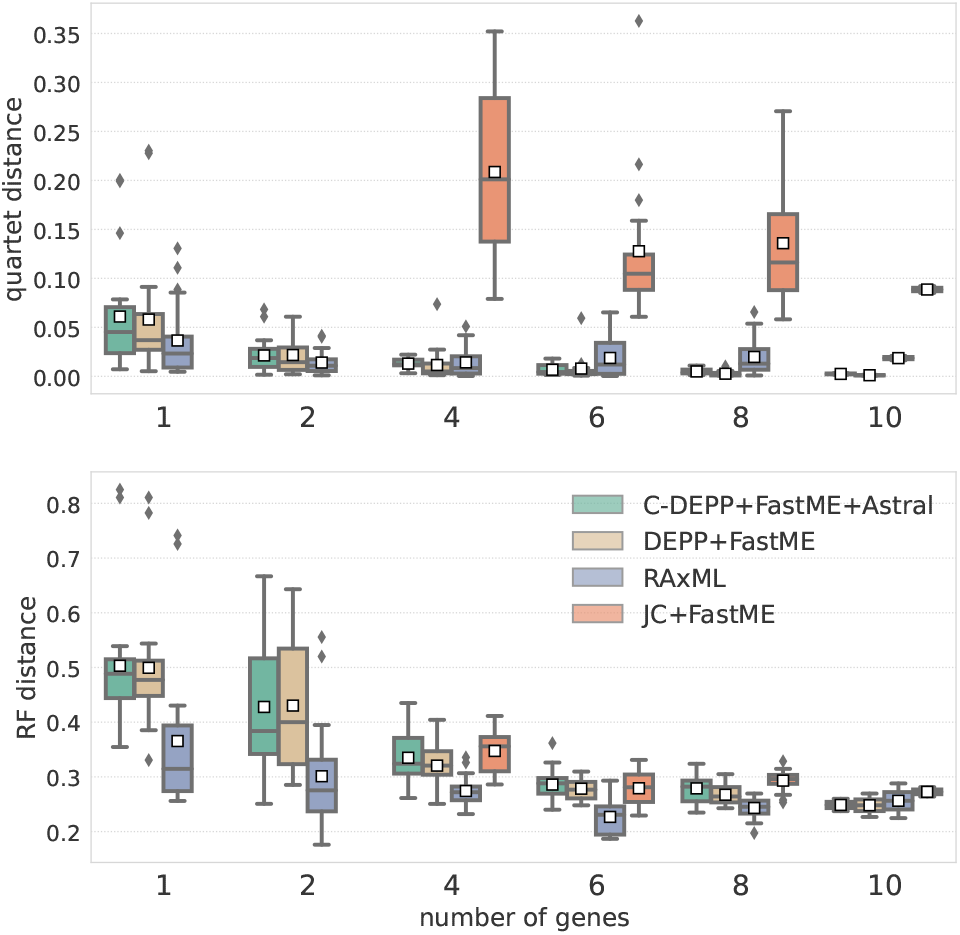
Tree update results on the WoL dataset. We show both quartet and RF distances, only restricted to query taxa, as the number of genes increases. FastME fails to run in some cases with one or two genes because specific pairs of sequences occasionally have no overlapping non-gap sites.

### Real world application

Recently, McDonald et al. (2022) inferred a reference tree combining ≈ 16, 000 genomes and 321,210 16S full-length sequences to produce the second version of the popular Greengenes database. The next goal of that project was to place all 23,113,447 short V4 16S rRNA Deblur v1.1.0 8 (Amir et al., 2017) amplicon sequence variants from Qiita (Gonzalez et al., 2018) (retrieved Dec. 14, 2021) on this tree. Since the backbone tree is a mixed species/gene tree, DEPP, which can learn to place on any tree, was the appropriate placement method. However, because the backbone tree was more than an order of magnitude larger than what DEPP could handle, we had to develop the C-DEPP approach studied here (an earlier version akin to TreePart+Class-Multi). McDonald et al. (2022) extensively reports on the results of that analyses showing improved taxonomic classification and consistency across data types in using GG2. Here, instead of repeating those results, we focus on the impact of C-DEPP. We study the impact of enabling DEPP to handle a backbone tree of size *>* 3 × 10^5^ by showing that using the original backbone of size 10^4^ would lead to more basal placements (Figure 6). The reduced terminal branch lengths indicate that a larger backbone tree can provide more precise phylogenetic placements and enable a better understanding of the microbiome compositions. Comparing the running time, placement using C-DEPP onto the backbone tree with more than 330,000 leaves requires roughly the same time as placement onto the WoL tree (with 10,575 leaves) using DEPP, further demonstrating the impressive scalability of C-DEPP.

**Fig. 6.**
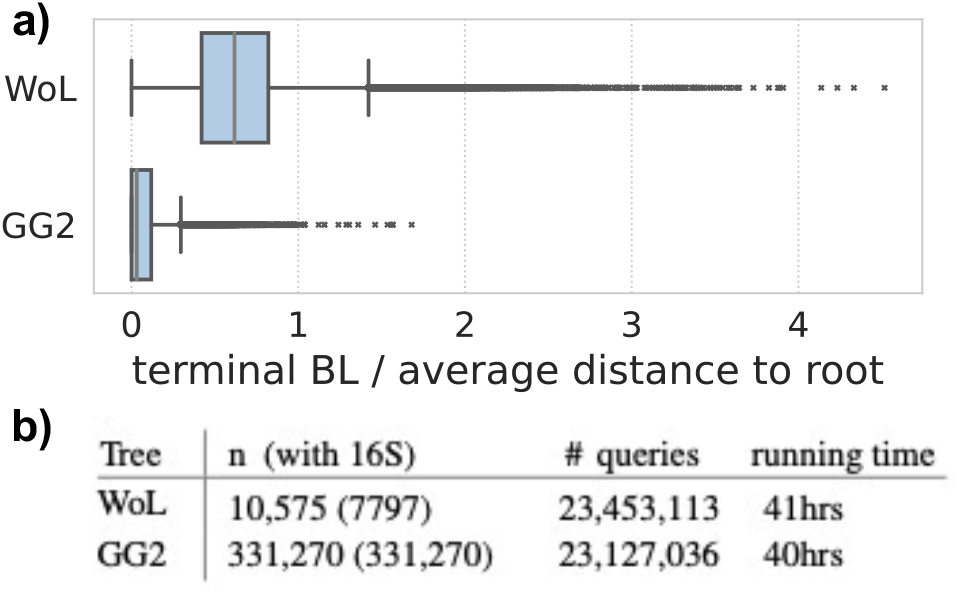
Impact of using C-DEPP on GG2 reference set with the THDMI dataset. a) Terminal branch length of the leaves in WoL reference tree and GG2 reference tree. b) Statistics of placement onto WoL reference tree and GG2 reference tree. Notice that the running time includes the time for sequence alignment and use a machine with 128 cores.

## Discussions

We presented a method called C-DEPP that enables the distance-based machine-learning placement method DEPP to scale to ultra-large datasets. C-DEPP is not more accurate than normal DEPP but is far more scalable as the backbone size grows. Unlike DEPP, which requires quadratic memory and running time, the time and memory of C-DEPP grow quasi-linearly with the backbone size. As more microbial genomes become available, the need for sub-quadratic methods becomes increasingly common. The C-DEPP method is already being used by other projects (McDonald et al., 2022) and fills an important practical gap; the most compelling reason for the need for C-DEPP is the practical applications it enables. The alternative distance-based methods (e.g., JC model within APPLES-II) are not accurate enough while the maximum-likelihood-based methods are too slow. C-DEPP is as salable (or perhaps more salable) than simple distance-based methods but comes close to maximum-likelihood in terms of accuracy.

In designing C-DEPP, we used common divide-and-conquer techniques but with several important twists. Among these, two stand out: 1) Phylogenetic-based division works only if partitions are augmented by representatives from other groups. This notion relates to the need for the sequence embedders to “see” sufficiently diverse examples to learn generalizable models; as such, this benefit may be particular to machine learning approaches and may not extend to standard phylogenetic methods. This reason for creating over-lapping subsets is very different from some existing work (e.g., Nelesen et al., 2012), which create overlapping subsets to allow merging of trees. 2) It is beneficial (albeit, mildly) to restrict distance calculation for each query to models more likely to have generated it. However, this benefit is enjoyed *only* if a two-layer classifier is used for judging the likelihood. A biological insight underpins this two-layer design. The main cause of gene tree discordance on microbial datasets is HGT. In the presence of HGT, classifying a gene into groups defined by the species tree can be misleading because the relevant part of the tree according to the gene tree may be different from the species tree. By making classification groups small, we give the classifier a chance to find clades in the species tree corresponding to the HGT origin in a specific gene.

Both the method and our analyses can improve in the future in several ways. In particular, for the tree update pipeline, we have made several choices without exploring alternatives. For example, we summarize genes by computing the median distance across genes; the alternative is to simply compute an updated tree per gene and let ASTRAL combine them. We also did not force the backbone trees to be fixed. It is possible to impose such constraints using ASTRAL (Rabiee and Mirarab, 2020a), and the constraints may improve the accuracy of the query relationship. We leave the exploration of these alternatives to the future. The most accurate method, SCAMPP, was slow in our analyses. A new version called Batch-SCAMPP (Wedell et al., 2022b) promises much faster running times. In preliminary tests, we saw some improvements in speed but not enough to bridge the gap with C-DEPP. Batch-SCAMPP takes between 2.5 and 9 hours on the 64k dataset, depending on the level of HGT in a gene, compared to 12 hours for normal SCAMPP. In contrast, C-DEPP takes around 30 minutes. Finally, while in this paper we scaled to trees with up to 330,000 leaves, larger trees have been built more recently. In our future applications of C-DEPP, we plan to use it to place queries on trees with a million leaves.

## Supplementary material: C-DEPP

**Fig. S7.**
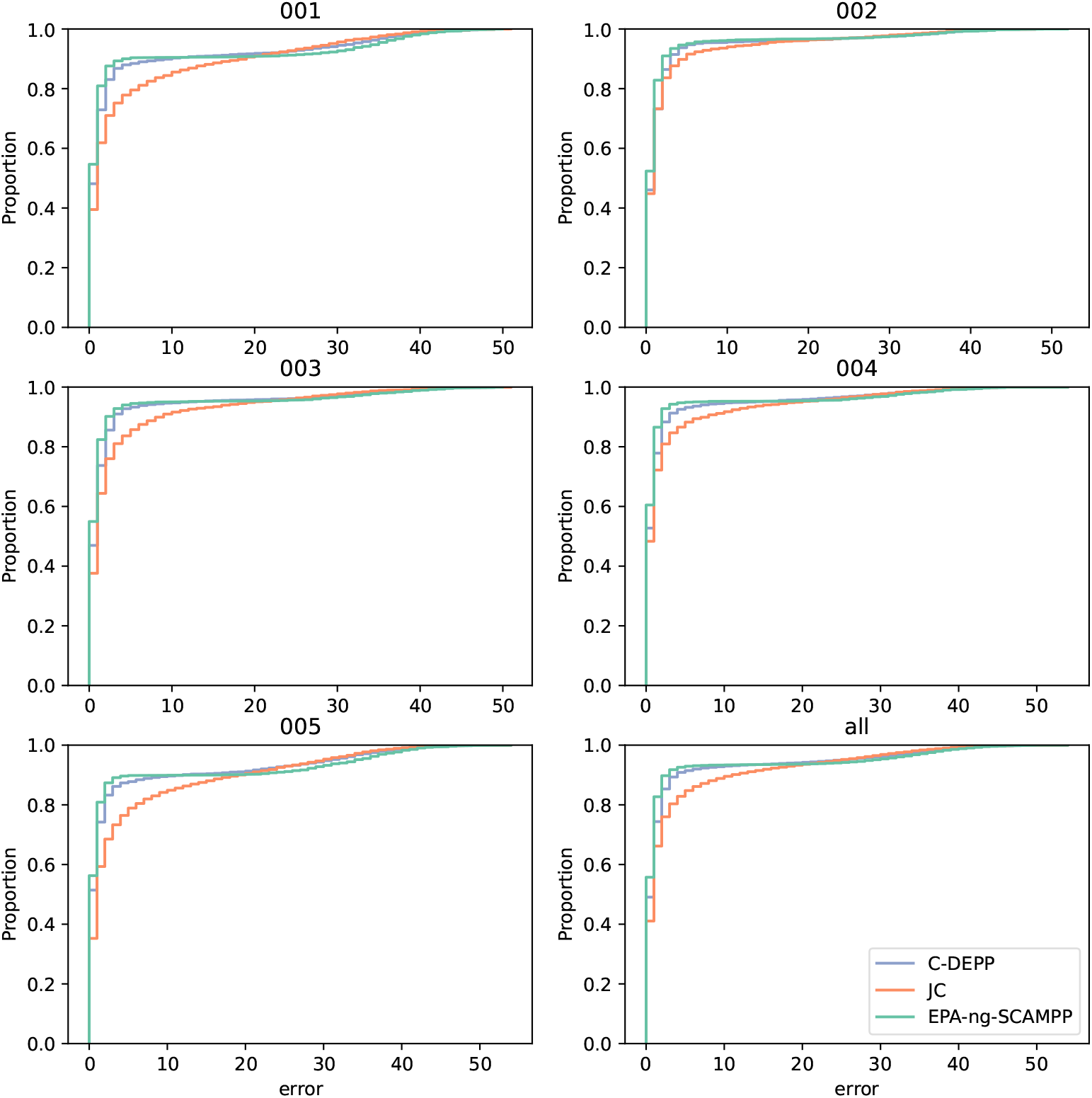
empirical cumulative distribution functions (ECDF) of errors in HGT-Sim (64K) dataset

